# The genetic landscape for amyloid beta fibril nucleation accurately discriminates familial Alzheimer’s disease mutations

**DOI:** 10.1101/2020.09.22.308429

**Authors:** Mireia Seuma, Andre Faure, Marta Badia, Ben Lehner, Benedetta Bolognesi

## Abstract

Amyloid fibrils are associated with many human diseases but how mutations alter the propensity of proteins to form fibrils has not been comprehensively investigated and is not well understood. Alzheimer’s Disease (AD) is the most common form of dementia with amyloid plaques of the amyloid beta (Aß) peptide a pathological hallmark of the disease. Mutations in Aß also cause familial forms of AD (fAD). Here we use deep mutational scanning to quantify the effects of >14,000 mutations on the aggregation of Aß. The resulting genetic landscape reveals fundamental mechanistic insights into fibril nucleation, including the importance of charge and gatekeeper residues in the disordered region outside of the amyloid core in preventing nucleation. Strikingly, unlike computational predictors and previous measurements, the *in vivo* nucleation scores accurately identify all known dominant fAD mutations, validating this simple cell-based assay as highly relevant to the human genetic disease and suggesting accelerated fibril nucleation is the ultimate cause of fAD. Our results provide the first comprehensive map of how mutations alter the formation of any amyloid fibril and a validated resource for the interpretation of genetic variation in Aß.

**Highlights:**

- First comprehensive map of how mutations alter the propensity of a protein to form amyloid fibrils.
- Charge and gatekeeper residues in the disordered N-terminus of amyloid beta prevent fibril nucleation.
- Rates of nucleation in a cell-based assay accurately identify the mutations that cause dominant familial Alzheimer’s disease.
- The combination of deep mutational scanning and human genetics provides a general strategy to quantify the disease-relevance of *in vitro* and *in vivo* assays.

## Introduction

Amyloid plaques consisting of the amyloid beta (Aß) peptide are a pathological hallmark of Alzheimer’s disease (AD), the most common cause of dementia and a leading global cause of morbidity with very high societal and economic impact (Ballard et al. 2011; “WHO Dementia: A Public Health Priority,” n.d.). Although most cases of AD are sporadic and of uncertain cause, rare familial forms of the disease also exist (Campion et al. 1999). These inherited forms of dementia typically have earlier onset and are caused by high penetrance mutations in multiple loci, including in the amyloid precursor protein (APP) gene, which encodes the protein from which Aß is derived by proteolytic cleavage (O’Brien and Wong 2011). The two most abundant isoforms of Aß are 42 and 40 amino acids (aa) in length, with the longer Aß42 peptide associated with increased aggregation *in vitro* and cellular toxicity (Meisl et al. 2014; Sandberg et al. 2010). The amyloid state is a thermodynamically low energy state but, both *in vitro* and *in vivo*, the spontaneous formation of amyloids is normally very slow because of the high kinetic barrier of fibril nucleation (Knowles, Vendruscolo, and Dobson 2014). The process of nucleation generates oligomeric Aß species that have been hypothesized to be particularly toxic to cells and that then grow into fibrils (Michaels et al. 2020; Bolognesi et al. 2010; Cleary et al. 2005).

Fourteen different mutations in the Aß42 peptide have been reported to cause familial Alzheimer’s disease (fAD), with all but two having a dominant pattern of inheritance (Weggen and Beher 2012; Van Cauwenberghe, Van Broeckhoven, and Sleegers 2016). However, it is not clear why these particular mutations cause fAD (Weggen and Beher 2012; Van Cauwenberghe, Van Broeckhoven, and Sleegers 2016), and these 14 mutations represent only 3.7% of the possible 378 single nucleotide changes that can occur in Aß. As for nearly all disease genes, therefore, the molecular mechanism by which mutations cause the disease remains unclear and the vast majority of possible mutations in Aß are variants of uncertain significance (VUS). This makes the clinical interpretation of genetic variation in this locus a difficult challenge (Starita et al. 2017; Gelman et al. 2019). Moreover, given the human mutation rate and population size, it is likely that nearly all of these possible variants in Aß actually exist in at least one individual currently alive on the planet (Conrad et al. 2011). A comprehensive map of how all possible mutations affect the formation of Aß amyloids and how these changes relate to the human disease is therefore urgently needed.

More generally, amyloid fibrils are associated with many different human diseases (Knowles, Vendruscolo, and Dobson 2014), but how mutations alter the propensity of proteins to aggregate into amyloid fibrils is not well understood and there has been no large-scale analysis of the effects of mutations on the formation of any amyloid fibril. Here we address this shortcoming by quantifying the rate of fibril formation for >14,000 variants of Aß. This provides the first comprehensive map of how mutations alter the propensity of any protein to form amyloid fibrils. The resulting data provide numerous mechanistic insights into the process of Aß fibril nucleation. Moreover, they also accurately classify all the known dominant fAD mutations, validating the clinical relevance of a simple cell-based model and providing a comprehensive resource for the interpretation of clinical genetic data.

## Results

### Deep Mutagenesis of Aß

To globally quantify the impact of mutations on the nucleation of Aß fibrils, we used an *in vivo* selection assay in which the nucleation of Aß is rate limiting for the aggregation of a second amyloid, the yeast prion [*PSI+*] encoded by the *sup35* gene (Chandramowlishwaran et al. 2018). Aggregation of Sup35p causes read-through of UGA stop codons, allowing growth based-selection using an auxotrophic marker containing a premature termination codon (Figure 1A). We generated a library containing all possible single nucleotide variants of Aß42 fused to the nucleation (N) domain of Sup35p and quantified the effect of mutations on the rate of nucleation in triplicate by selection and deep sequencing (see Methods). The selection assay was highly reproducible, with enrichment scores for aa substitutions strongly correlated between replicates (Figure 1B, figure supplement 1A).

**Figure 1.**
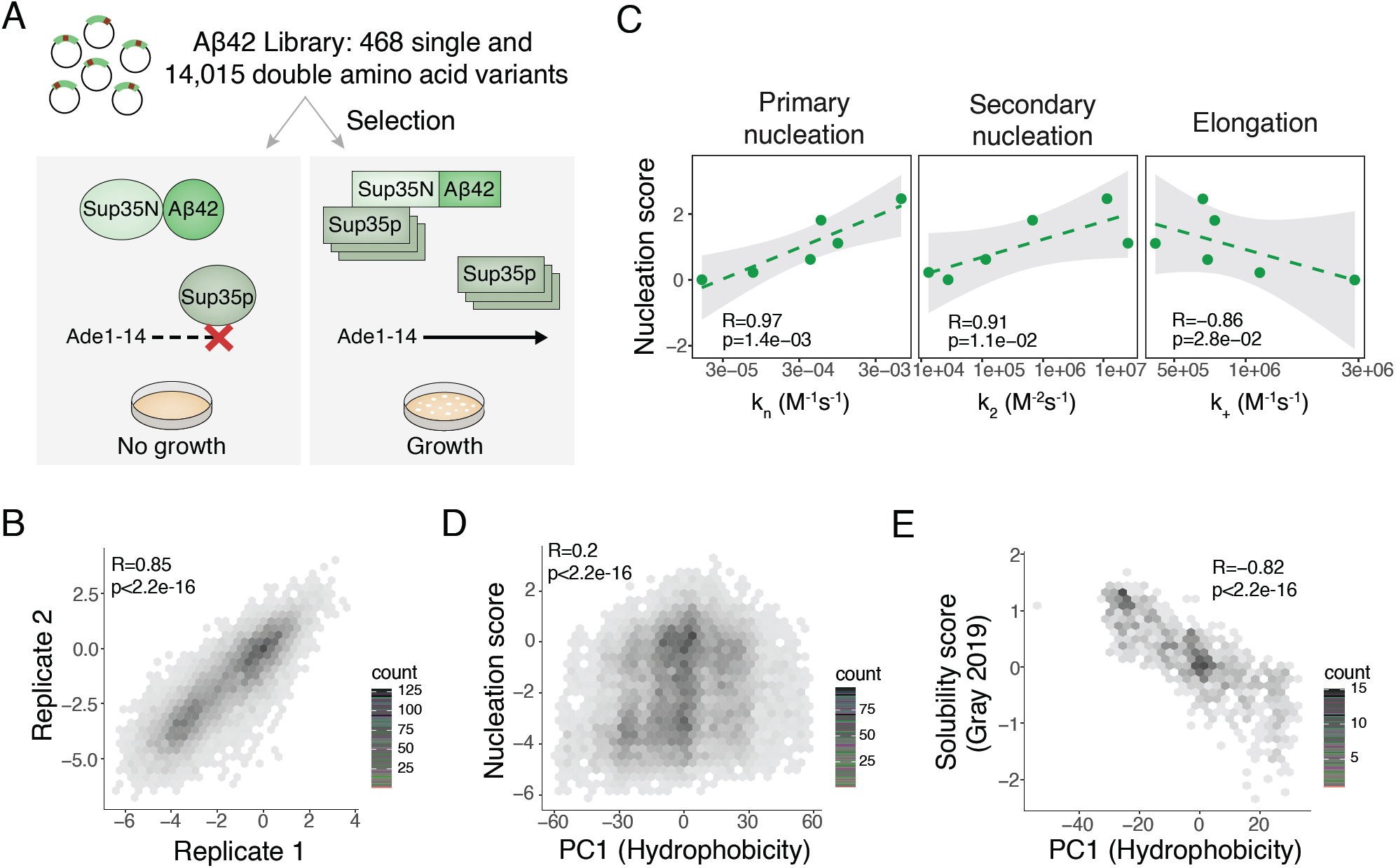
Deep mutagenesis of Aß nucleation. **A** *in vivo* Aß selection assay. Aß fused to the Sup35N domain seeds aggregation of endogenous Sup35p causing a readthrough of a premature UGA in the *Ade1-14* reporter, allowing the cells to grow in medium lacking adenine. **B** Correlation of nucleation scores for biological replicates 1 and 2 for single and double aa mutants. Pearson correlation coefficient and p-value are indicated (figure supplement 1A) n=10,157 genotypes **C** Correlation of nucleation scores with *in vitro* primary and secondary nucleation and elongation rate constants (Yang et al. 2018). Weighted Pearson correlation coefficient and p-value are indicated. **D** Nucleation scores as a function of principal component 1 (PC1) aa property changes (Bolognesi et al. 2019) for single and double aa mutants (n=14,483 genotypes). Weighted Pearson correlation coefficient and p-value are indicated. **E** Solubility scores (Gray et al. 2019) as a function of PC1 changes (Bolognesi et al. 2019) for n=895 single and double mutants. Pearson correlation coefficient and p-value are indicated.

### *in vivo* nucleation scores are highly correlated with *in vitro* rates of amyloid nucleation

Comparing our *in vivo* enrichment scores to previously quantified *in vitro* measures of Aß aggregation validated the assay and revealed that the *in vivo* scores correlate extremely well with the rate of Aß nucleation (Yang et al. 2018; Törnquist et al. 2018) (Figure 1C, figure supplement 1B). We henceforth refer to the *in vivo* enrichment scores as ‘nucleation scores’.

### Two mechanisms of *in vivo* Aß aggregation

A prior deep mutational scan quantified the effects of mutations on the abundance of Aß fused to an enzymatic reporter (Gray et al. 2019). These ‘solubility scores’ do not predict the effects of mutations on Aß nucleation (figure supplement 1C). Previously we identified a principal component of aa properties (PC1, related to changes in hydrophobicity) that predicts the aggregation and toxicity of the amyotrophic lateral sclerosis (ALS) protein TDP-43 (Bolognesi et al. 2019). PC1 is also not a good predictor of Aß nucleation (Figure 1D) but it does predict the previously reported changes in Aß solubility (Figure 1E), suggesting that Aß is aggregating by a similar process to TDP-43 in this alternative selection assay (Gray et al. 2019) but by a different mechanism in the nucleation selection.

### Nucleation scores for 14,483 Aß variants

The distribution of mutational effects for Aß nucleation has a strong bias towards reduced nucleation, with 56% of single aa substitutions reducing nucleation but only 16% increasing it (Z-test, false discovery rate, FDR=0.1, Figure 2A). Moreover, mutations that decrease nucleation in our dataset typically have a larger effect than those that increase it, with many mutations reducing nucleation to the background rate observed for Aß variants containing premature termination codons (Figure 2A).

**Figure 2.**
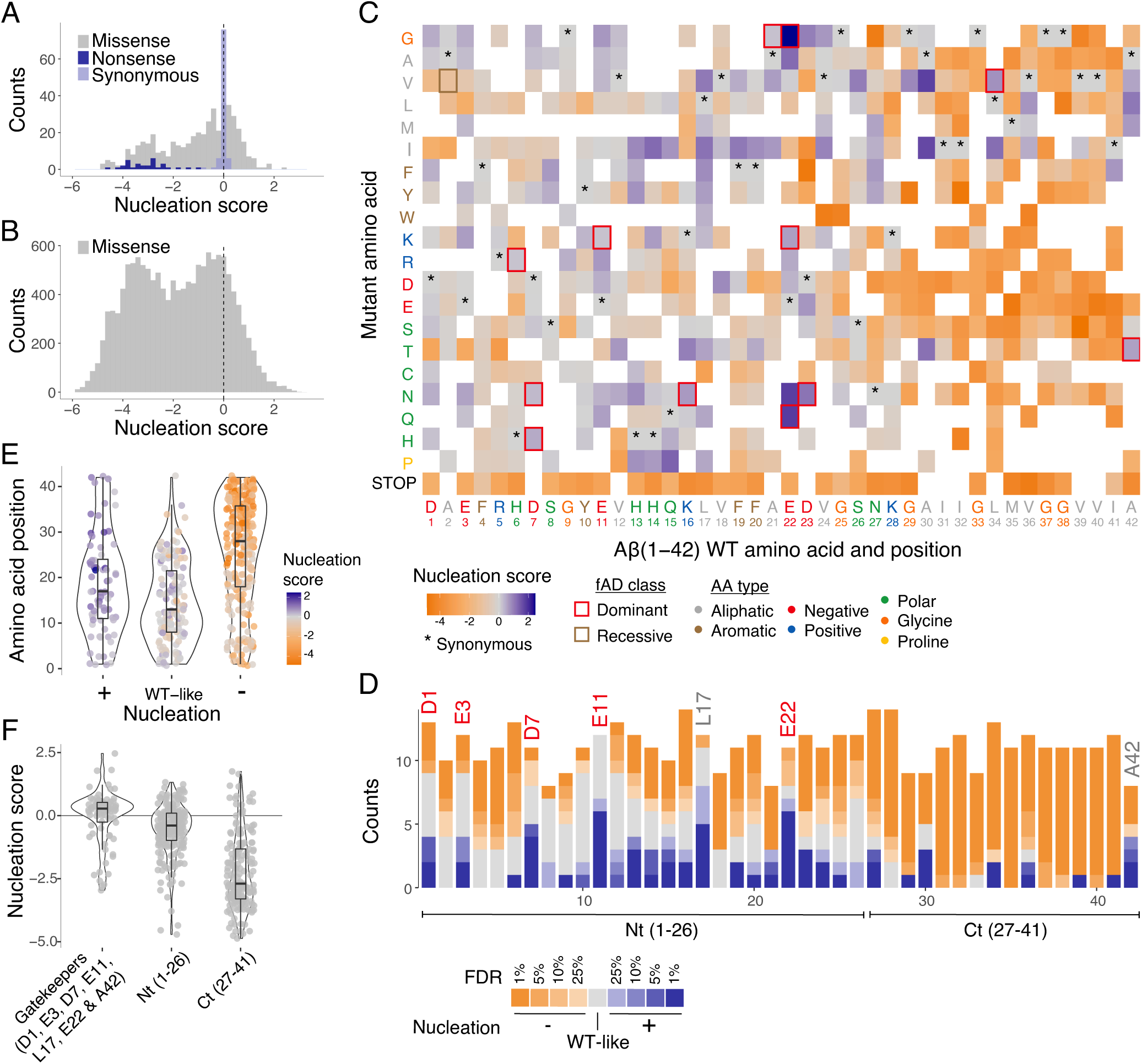
Modular organisation of mutational effects in Aß. **A and B** Nucleation scores distribution for single (**A**) and double (**B**) aa mutants. n=468 (Missense), n=31 (Nonsense), n=90 (Synonymous) for singles, and n=14,015 (Missense) for doubles. Vertical dashed line indicates WT score (0). **C** Heatmap of nucleation scores for single aa mutants. The WT aa and position are indicated in the x-axis and the mutant aa is indicated on the y-axis, both coloured by aa class. Variants not present in the library are represented in white. Synonymous mutants are indicated with ‘*’ and fAD mutants with a box, coloured by fAD class. **D** Number of variants significantly increasing (blue) and decreasing (orange) nucleation at different FDRs. Gatekeeper positions (D1, E3, D7, E11, L17, E22 and A42) are indicated on top of the corresponding bar and coloured on the basis of aa type. The N-terminal and C-terminal definitions are indicated on the x-axis. Gatekeeper positions are excluded from the N-terminal and C-terminal classes. **E** aa position distributions for variants that increase (+) decrease (−) or have no effect on nucleation (WT-like) (FDR<0.1). **F** Nucleation score distributions for the three clusters of positions defined on the basis of nucleation: Nt (1-26), Ct (27-41) and gatekeeper positions (clusters are mutually exclusive). Horizontal line indicates WT nucleation score (0). Nonsense (stop) mutants were only included in **A** and **C**. Boxplots represent median values and the lower and upper hinges correspond to the 25th and 75th percentiles respectively. Whiskers extend from the hinge to the largest value no further than 1.5*IQR (interquartile range). Outliers are plotted individually or omitted when the boxplot is plotted together with individual data points or a violin plot.

In addition to covering all aa changes obtainable through single nt mutations, our mutagenesis library was designed to contain a substantial fraction of double mutants. In total, we quantified the impact of 14,015 double aa variants of Aß. Double mutants were even more likely to reduce nucleation, with 63% decreasing and only 5.5% increasing nucleation (Z-test, FDR=0.1, Figure 2B). Therefore, mutations more frequently decrease rather than increase Aß nucleation.

### Aß has a modular mutational landscape

Inspecting the heat map of mutational effects for aa changes at all positions in Aß reveals strong biases in the locations of mutations that increase and decrease nucleation (Figure 2C and D, figure supplement 2A). Mutations that decrease nucleation are highly enriched in the C-terminus of Aß whereas mutations that increase nucleation are enriched in the N-terminus (Figure 2E). Indeed, >84% of mutations in the C-terminus (residues 27-42) reduce nucleation and only 9.6% increase it (FDR=0.1). In contrast, the effects of mutations are smaller (Figure 2F) and also more balanced in the first 26 aa of the peptide, with 38.6% decreasing and 20% increasing nucleation (FDR=0.1).

These differences in the direction and strength of mutational effects between the N- and C-terminal regions of Aß suggest a modular organization of the peptide. This modularity is also reflected in the primary sequence of Aß, which has a hydrophobic C-terminus and a more polar and charged N-terminus (8 out of 9 charged residues in Aß are found before residue 24 and the peptide consists entirely of hydrophobic residues from position 29) (Figure 2C, 3A). Consistent with this modular organisation, mutations in the few hydrophobic residues in the N-terminus have effects that are more similar to mutations in polar residues in the N-terminus rather than in hydrophobic residues in the C-terminus. Similarly, mutations in the most C-terminal charged residue (K28) frequently strongly reduce nucleation, just as they do in the adjacent hydrophobic positions (figure supplement 2B).

### Gatekeeper residues act as anti-nucleators

Considering the entire Aß peptide, there are only seven positions in which mutations are not more likely to decrease rather than increase nucleation (FDR=0.1, Figure 2D). Strikingly, these positions, which we refer to as ‘gatekeepers’ of nucleation (Rousseau, Serrano, and Schymkowitz 2006; Pedersen, Christensen, and Otzen 2004), include five of the six negatively charged residues in Aß. The sixth gatekeeper is an unusual hydrophobic residue in the N-terminus, L17, where 7 mutations increase nucleation and only one decreases it (FDR=0.1, Figure 2D). The final aa of the peptide, A42, also has an unusual distribution of mutational effects that is different to the rest of the C-terminus, with four mutations increasing and three mutations decreasing nucleation (FDR=0.1, Figure 2D).

Taken together, on the basis of mutational effects, we therefore distinguish the following mutually exclusive positions in Aß: the C-terminus (aa 27-41) where the majority of mutations strongly decrease nucleation, the N-terminus (aa 1-26) where mutations have smaller and more balanced effects, and seven gatekeeper residues (D1, E3, D7, E11, D22, L17, A42) where mutations frequently increase nucleation. We consider each of these classes below.

### Mutations in the N- and C-terminal regions

Mutations in the C-terminus nearly all decrease nucleation (Figure 3A). This is consistent with the C-terminus forming part of the tightly-packed amyloid core of all known structural polymorphs of both Aß42 (Colvin et al. 2016; Meier, Riek, and Böckmann 2017; Wälti et al. 2016; Xiao et al. 2015; Gremer et al. 2017; Lührs et al. 2005; Schmidt et al. 2015) and Aß40 (Kollmer et al. 2019; Lu et al. 2013; Qiang et al. 2012; Sgourakis, Yau, and Qiang 2015; Paravastu et al. 2008; Schütz et al. 2015). Mutations to polar and charged residues in this region nearly all decrease nucleation, but so too do most changes to other hydrophobic residues (Figure 3B), suggesting specific side chain packing in this region is important for nucleation. The relative effects of different mutations are only partially captured by changes in hydrophobicity (Figure 3F; Pearson correlation coefficient, R=0.45) and by predictors of aggregation potential (figure supplement 3A). Only a few mutations in this region increase nucleation: substitutions to isoleucine at positions 30, 34 and 39; mutations to valine at positions 29, 30 and 34; a change to threonine at position 30; changes to leucine and methionine at 36, and mutation to phenylalanine at position 41 (FDR=0.1).

**Figure 3.**
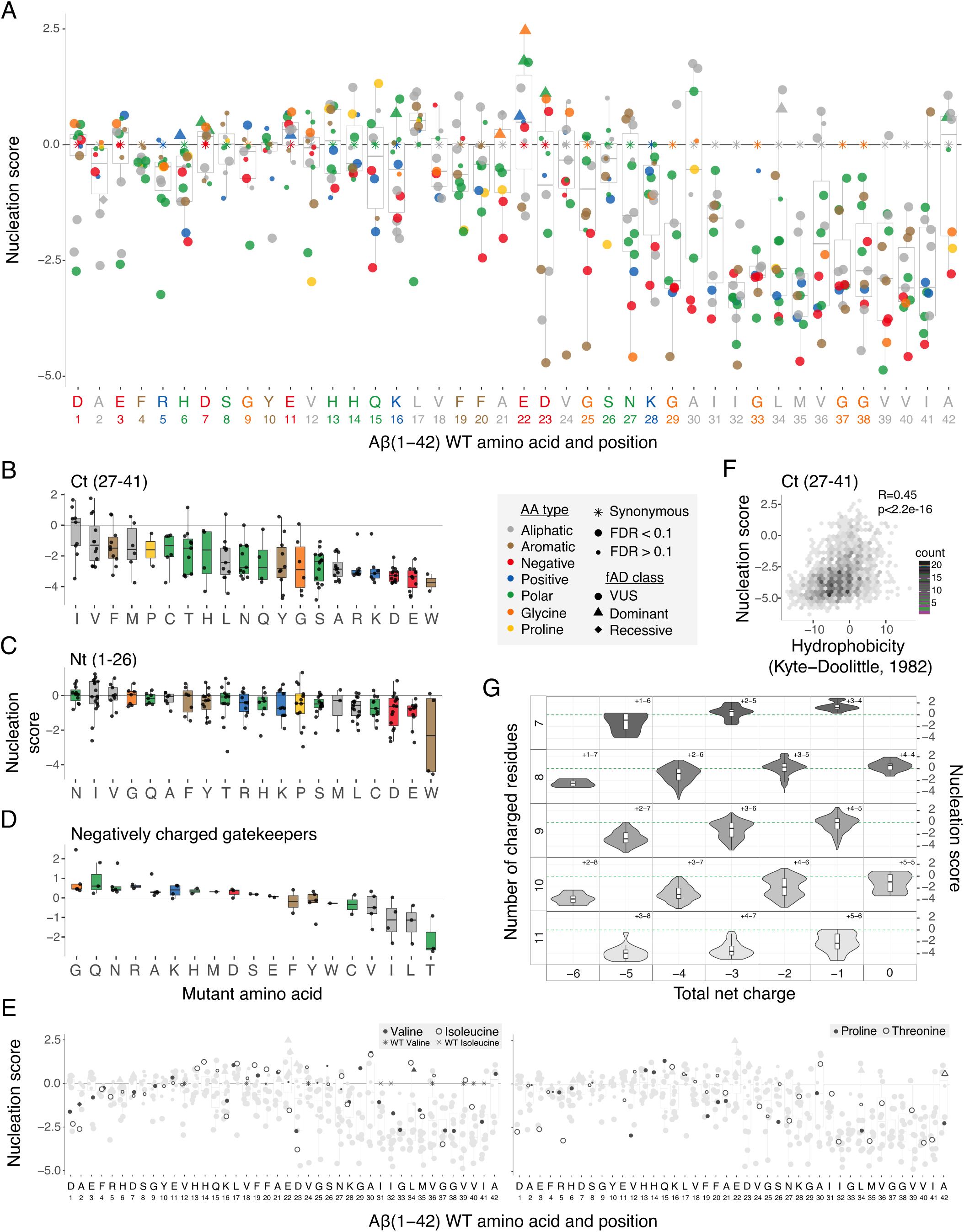
Determinants of Aß nucleation. **A** Effect of single aa mutants on nucleation for each Aß position. The WT aa and position are indicated on the x-axis and coloured on the basis of aa type. The horizontal line indicates the WT nucleation score (0). **B to D** Effect of each mutant aa on nucleation for the Ct (27-41) (**B**), the Nt (1-26) (**C**) and the negatively charged gatekeeper positions (D1, E3, D7, E11 and E22) (**D**). The 3 position clusters are mutually exclusive. Colour code indicates aa type.The horizontal line is set at the WT nucleation score (0). **E** Effect on nucleation for single aa mutations to proline, threonine, valine and isoleucine. Mutations to other aa are indicated in grey. The horizontal line indicates WT nucleation score (0). Point size and shape indicate FDR and fAD class respectively (see legend). **F** Nucleation scores as a function of hydrophobicity changes (Kyte and Doolittle 1982) for single and double aa mutants in the Ct (27-41) cluster. Only double mutants with both mutations in the indicated position-range were used. Weighted Pearson correlation coefficient and p-value are indicated. **G** Nucleation score distributions arranged by the number of charged residues (y-axis) and the total net charge (x-axis) for single and double aa mutants in the full peptide (1-42). Only polar, charged and glycine aa types were taken into account, for both WT and mutant residues. Colour gradient indicates the total number of charged residues. Numbers inside each cell indicate the number of positive and negative residues. The horizontal line indicates WT nucleation score (0). Boxplots represent median values and the lower and upper hinges correspond to the 25th and 75th percentiles respectively. Whiskers extend from the hinge to the largest value no further than 1.5*IQR (interquartile range). Outliers are plotted individually or omitted when the boxplot is plotted together with individual data points or a violin plot.

Mutations in the N-terminus of Aß have a more balanced effect on nucleation, and these effects are not well predicted by either hydrophobicity or predictors of aggregation potential (figure supplement 3B, C and D). The effects of introducing particular amino acids are, however, biased, with the introduction of N, I, and V most likely to increase nucleation (Figure 3C and figure supplement 4). As at the C-terminus, the introduction of negative charged residues typically strongly reduces nucleation (Figure 3B and C). However, in contrast to what is observed in the C-terminus (Figure 3B), the effects of introducing positive charge are less severe (Figure 3C). Interestingly, the effects of mutations to proline, isoleucine, valine and threonine in the N-terminus depend on the position in which they are made: mutations in the first 12 residues typically decrease nucleation whereas mutations in the next four to nine residues increase nucleation (Figure 3E). The conformational rigidity of P and the beta-branched side chains of I, V and T that disfavour helix formation suggest that disruption of a secondary structure in this region may favour nucleation.

### The role of charge in limiting Aß nucleation

At five of six negatively charged positions in Aß, mutations frequently increase nucleation (Figure 2D and 3A). Moreover, the introduction of negative charge at other positions strongly decreases nucleation (Figure 3A), suggesting that negatively charged residues act as gatekeepers (Pedersen, Christensen, and Otzen 2004; Rousseau, Serrano, and Schymkowitz 2006) to limit nucleation (Figure 3D and figure supplement 3C). In contrast, mutations in the three positively charged residues (R5, K16, K28) mostly decrease nucleation (Figure 2D). Mutating the negatively charged gatekeepers to the polar amino acids Q and N, to positively charged residues (R and K), or to small side chains (G and A) increases nucleation (Figure 3D). Conversely, mutating the same positions to hydrophobic residues typically reduces nucleation (Figure 3D). This is consistent with a model in which the negative charge at these positions acts to limit nucleation, but that the overall polar and unstructured nature of the N-terminus must be maintained for effective nucleation.

To further investigate the role of charge in controlling Aß nucleation, we extended our analyses to the double mutants. Including double mutants allows the net charge of Aß to vary over a wider range and it also allows comparison of the nucleation of peptides with the same net charge but a different total number of charged residues (for example, a net charge of −3 can result from a negative/positive aa composition of 6/3, as in wild-type Aß, or compositions of 7/4, 5/2 etc). Considering all mutations between charged and polar residues or glycine reveals that, although reducing the net charge of the peptide from −3 progressively increases nucleation (Figure 3G), the total number of charged residues is also important: for a given net charge, nucleation is increased in peptides containing fewer charged residues of any sign (Figure 3G and figure supplement 3E and F). Thus, both the overall charge and the number of charged residues control the rate of Aß nucleation.

### Hydrophobic gatekeeper residues

In addition to the five negatively charged gatekeeper residues, mutations most frequently increase nucleation of Aß in two specific hydrophobic residues: L17 and A42 (Figure 2C and D). At position 17, changes to polar, aromatic and aliphatic amino acids all increase nucleation, as does the introduction of a positive charge and mutation to proline. Only a mutation to cysteine reduces nucleation (Figure 2C). This suggests a specific role for leucine at position 17 in limiting nucleation, perhaps as part of a nucleation-limiting secondary structure suggested by the mutational effects of P, I, V and T in this region (Figure 3E).

Finally, the distribution of mutational effects at position 42 differs to that in the rest of the hydrophobic C-terminus of Aß, with mutations most often increasing nucleation (Figure 2D, FDR=0.1). The mutations that increase nucleation are all to other aliphatic residues (Figure 2C and 3A). The distinction of position 42 is interesting because of the increased toxicity and aggregation propensity of Aß42 compared to the shorter Aß40 APP cleavage product (Meisl et al. 2014; Sandberg et al. 2010).

### Nucleation scores accurately discriminate fAD mutations

To investigate how nucleation in the cell-based assay relates to the human disease, we considered all the mutations in Aß known to cause familial AD (fAD). In total, there are 12 mutations in Aß reported to cause dominantly inherited fAD. These 12 known disease mutations are not well discriminated by commonly used computational variant effect predictors (Figure 4 and figure supplement 5A) or by computational predictors of protein aggregation and solubility (Figure 4 and figure supplement 5B). They are also poorly predicted by the previous deep mutational scan of Aß designed to quantify changes in protein solubility, suggesting the disease is unrelated to the biochemical process quantified in this assay (Gray et al. 2019) (figure supplement 5C).

**Figure 4.**
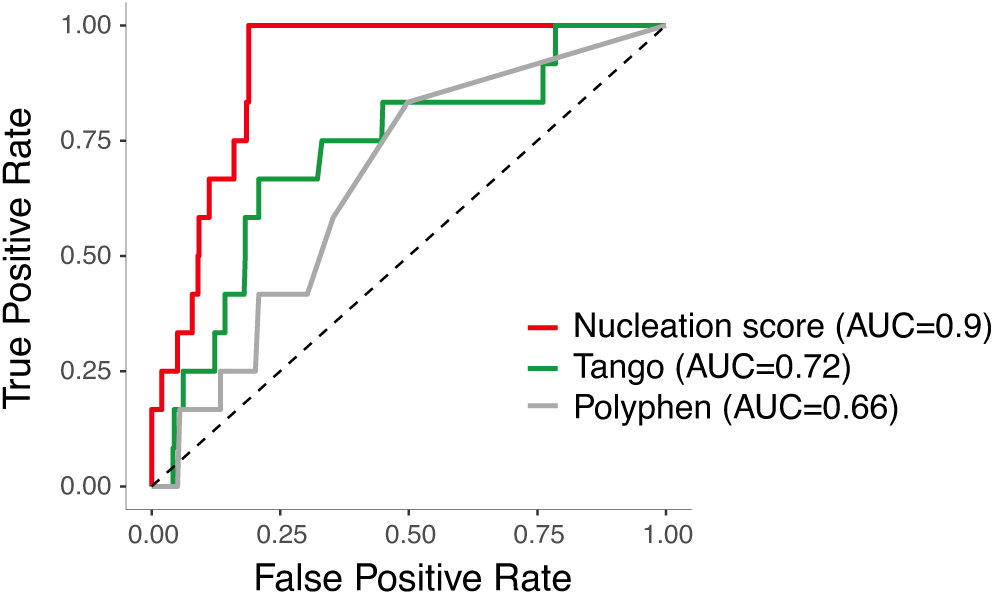
Aß nucleation accurately discriminates dominant fAD variants. ROC curves for 12 fAD mutants versus all other single aa mutants in the dataset. AUC values are indicated in the legend. Diagonal dashed line indicates the performance of a random classifier.

In contrast, the scores from our *in vivo* nucleation assay accurately classify the known dominant fAD mutations, with all 12 mutations increasing nucleation (Figure 4, area under the receiver operating curve, ROC-AUC=0.9, two-tailed Z-test, p<2.2e-16). This suggests the biochemical events occurring in this simple cell-based assay are highly relevant to the development of the human disease.

Consistent with the overall mutational landscape, the known fAD mutations are also enriched in the N-terminus of Aß (Figure 2C). In some positions the known fAD mutations are the only mutation or one of only a few mutations that can increase nucleation. For example, based on our data, H6R is likely to be the only dominant fAD mutation in position 6. However, in other positions there are multiple additional variants that increase nucleation as much as the known fAD mutation. At position 11, for example, there are five mutations with a nucleation score higher than the known E11K disease mutation (Figure 2C and D). Indeed, our data suggest there are likely to be many additional dominant fAD mutations beyond the 12 that have been reported to date (Supplementary file 1).

In addition to the 12 known dominant fAD mutations, two additional variants in Aß have been suggested to act recessively to cause fAD (Di Fede et al. 2009; Tomiyama et al. 2008). One of these variants is a codon deletion (E22Δ) and is not present in our library. The other variant, A2V, does not have a dominant effect on nucleation in our assay (Figure 2C), consistent with a recessive pattern of inheritance and a different mechanism of action. More generally, of the hundreds of amino acid changes possible in the peptide, our data prioritise 63 as candidate fAD variants (Supplementary file 1). 131 variants are likely to be benign, and 262 reduce Aß nucleation and so may even be protective. These include variants already reported in the gnomAD database of human genetic variation (figure supplement 5D).

## Discussion

Taken together, the data presented here provides the first comprehensive analysis of how mutations promote and prevent the aggregation of an amyloid, reveals a modular organisation for the impact of mutations on the nucleation of Aß, shows that the rate of nucleation in a cell-based assay accurately identifies mutations that cause dominant familial Alzheimer’s disease, and provides a resource for the future clinical interpretation of genetic variation in this locus.A majority of mutations in the C-terminal core of Aß disrupt nucleation, which is consistent with the formation of specific hydrophobic contacts in this region being required for nucleation. In contrast, mutations that increase nucleation are enriched in the polar N-terminus with mutations in negatively charged gatekeeper residues and the L17 gatekeeper being particularly likely to have a positive effect. Indeed, decreasing both the net charge of the peptide and the total number of charged residues increases nucleation. The mutational effects in residues immediately before position 17 are consistent with the formation of a structural element in this region interfering with nucleation. Little is known about the structure of Aß during fibril nucleation, but the results presented here are in general consistent with mature fibril structures of Aß where the C-terminal region of the peptide islocated in the amyloid core and the N-terminus isdisordered and solvent exposed (Figure 5, figure supplement 6 and 7).

**Figure 5.**
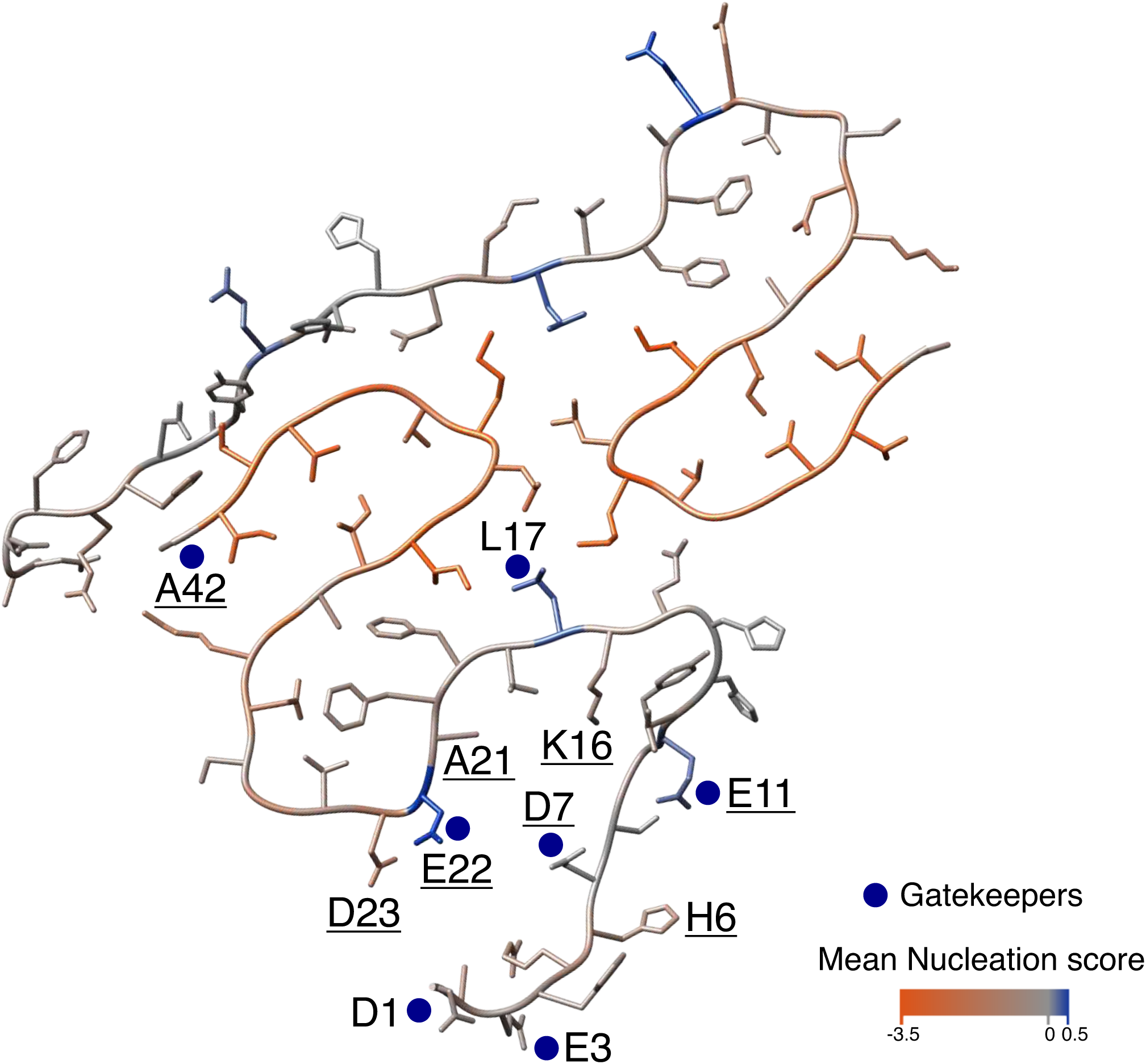
Mutational landscape of the Aß amyloid fibril. Average effect of mutations visualised on the cross-section of an Aß amyloid fibril (PDB accession 5KK3 (Colvin et al. 2016)). Nucleation gatekeeper residues and known fAD mutations positions are indicated by the WT aa identity on one of the two monomers; gatekeepers are indicated with blue dots and fAD are underlined. A single layer of the fibril is shown and the unstructured N-termini (aa 1 to 14) are shown with different random coil conformations for the two Aß monomers. See figure supplement 7 for alternative Aß42 amyloid polymorphs.

Our results also show that in the same type of cell (yeast), Aß aggregates in two different ways. One of these (quantified by the nucleation assay employed here) must be extremely similar to the aggregation that occurs in the human brain to cause familial Alzheimer’s disease. The other pathway of aggregation (quantified by the solubility assay (Gray et al. 2019)), however, is unrelated to the human disease. This second aggregation pathway is, at least to a large extent, driven by changes in hydrophobicity, similar to what we previously reported for the aggregation in yeast of the ALS protein, TDP-43 (Bolognesi et al. 2019).

More generally, our results highlight how the combination of deep mutational scanning and human genetics can be a general strategy to quantify the disease-relevance of biological assays. Many *in vitro* and *in vivo* assays are proposed as ‘disease models’ in biomedical research with their relevance often justified by how ‘physiological’ the assays seem or how well phenotypes observed in the model match those observed in the human disease. However, such criteria are largely subjective, and assays that seem relevant to a disease may actually turn out to be reporting on irrelevant biochemical events, resulting in the development of drugs that then fail in clinical trials.

Our study highlights an alternative approach, which is to use the complete set of known disease-causing mutations to quantify how well an assay recapitulates the biochemistry of a disease. Thus, although the yeast-based assay that we employed here might typically be dismissed as ‘non-physiological’, ‘artificial’ or ‘lacking many features important for a neurological disease’, unbiased massively parallel genetic analysis provides very strong evidence that it must be reporting on biochemical events that are extremely similar to – or the same as – those that occur to cause the human disease. Indeed, we would argue that this simple system is now better-validated as a model of fAD than any other, including animal models where the effects of only one or a few mutations (including control mutations) have ever been tested. Similarly-strong agreement between mutational effects in a cellular assay and the set of mutations already known to cause a disease is observed for other diseases (Starita et al. 2017; Gelman et al. 2019), suggesting the generality of this approach.

We suggest therefore that the combination of deep mutational scanning and human genetics provides a general strategy to quantify the disease-relevance of *in vitro* and *in vivo* assays. We encourage that deep mutagenesis should be employed early in discovery programmes to ‘genetically validate’ (or invalidate) the relevance of assays for particular diseases. The concordance between mutational effects in an assay and a disease is an unbiased metric that can be used to prioritise between different assays. Quantifying the ‘genetic agreement’ between an assay and a disease will help prevent time and resources being wasted on research that actually has little relevance to a disease.

Finally, the strikingly consistent effects of the dominant fAD mutations in our assay further strengthen the evidence that AD is actually a ‘nucleation disease’ ultimately caused by an increased rate of fibril nucleation (Aprile et al. 2017; Cohen et al. 2018; Knowles et al. 2009). Therapeutic strategies to prevent and treat AD should therefore focus on limiting this easily quantifiable biochemical process using genetically-validated assays such as the one employed here.

**figure supplement 1.**
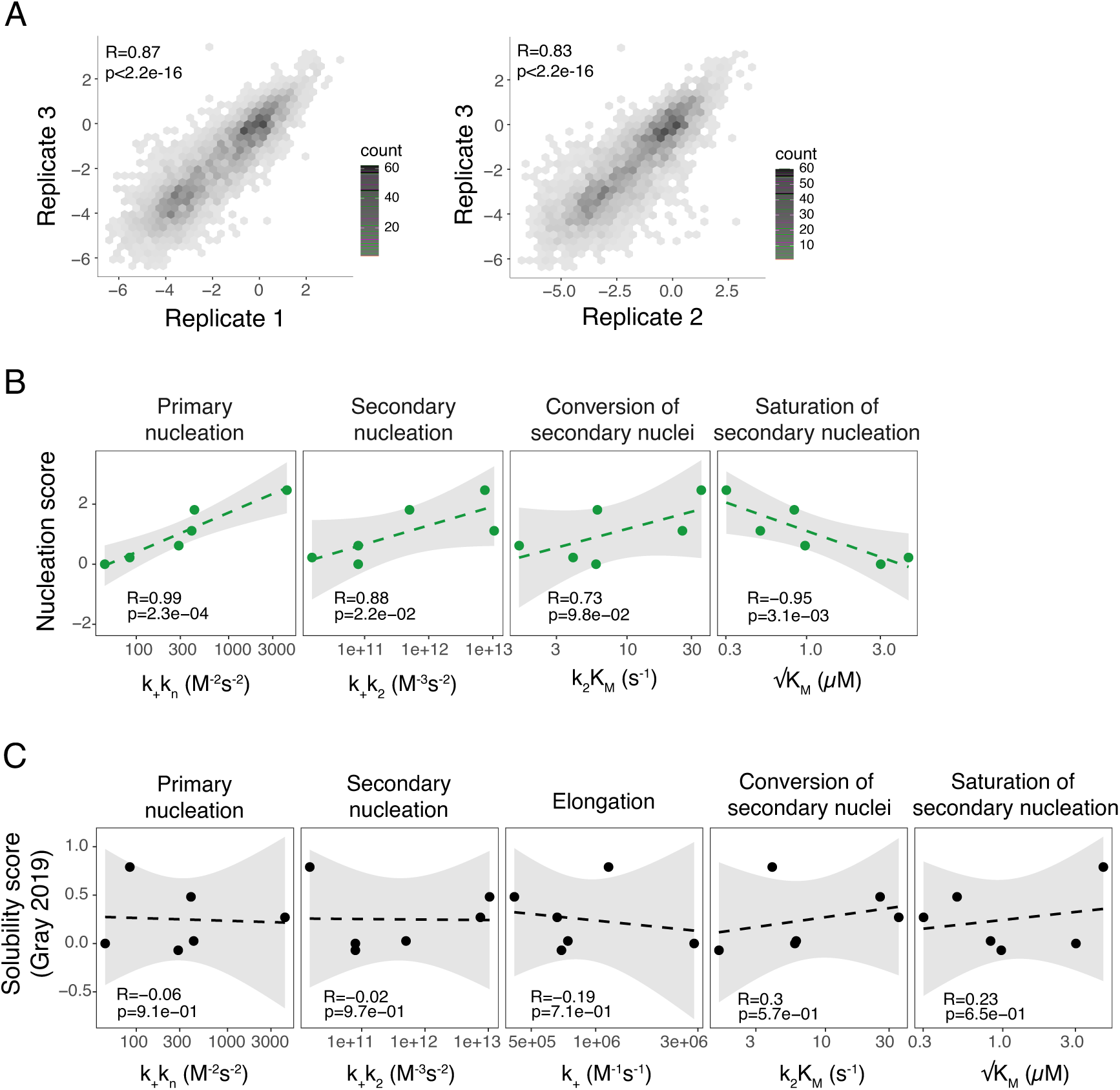
Reproducibility of the assay and correlation with *in vitro* fibril nucleation. **A** Correlation of nucleation scores for biological replicates 1 and 3 (n=4,124 genotypes) and 2 and 3 (n=4,093) for single and double aa mutants. Pearson correlation coefficient and p-value are indicated. **B** Correlation of nucleation scores with *in vitro* combined rate constants for primary nucleation, secondary nucleation, conversion of secondary nuclei and saturation of secondary nucleation (Yang et al. 2018). Weighted Pearson correlation coefficient and p-value are indicated. **C** Correlation of solubility scores (Yang et al. 2018) with *in vitro* combined rate constants for primary nucleation, secondary nucleation, elongation, conversion of secondary nuclei and saturation of secondary nucleation (Yang et al. 2018). Weighted Pearson correlation coefficient and p-value are indicated.

**figure supplement 2.**
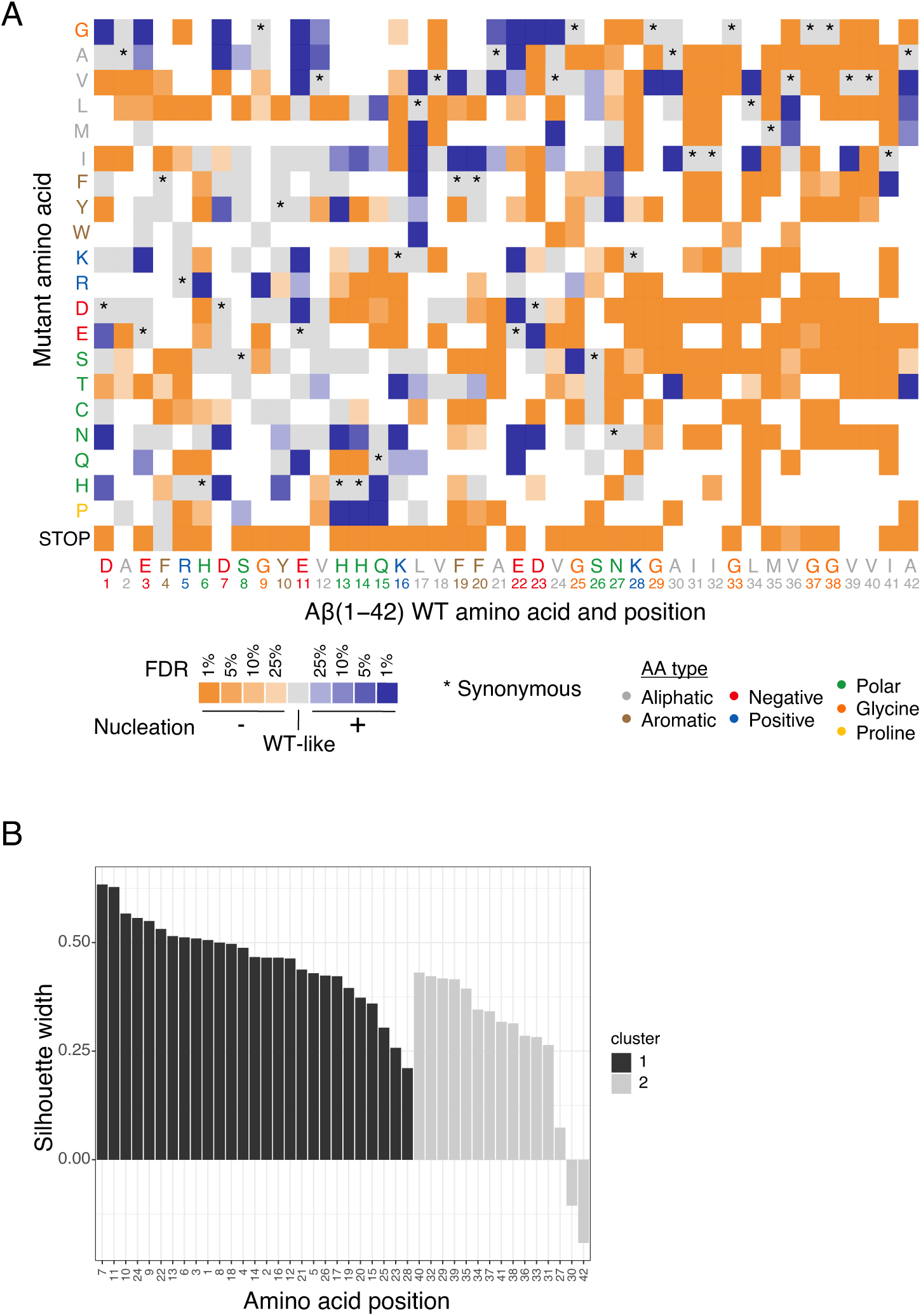
Mutational effects in Aß. **A** Heatmap of nucleation score categories for single aa variants at different FDRs (blue: increased nucleation, orange: decreased nucleation, grey: not different from WT). The WT aa and position are indicated on the x-axis and the mutant aa is indicated on the y-axis, both coloured on the basis of aa type class (see legend). Variants not present in the library are represented in white. Synonymous mutants are indicated with ‘*’. Nonsense mutations (stop) were included. **B** K-medoids clustering of single aa variant nucleation scores estimates by residue position. Silhouette widths for all positions with K=2.

**figure supplement 3.**
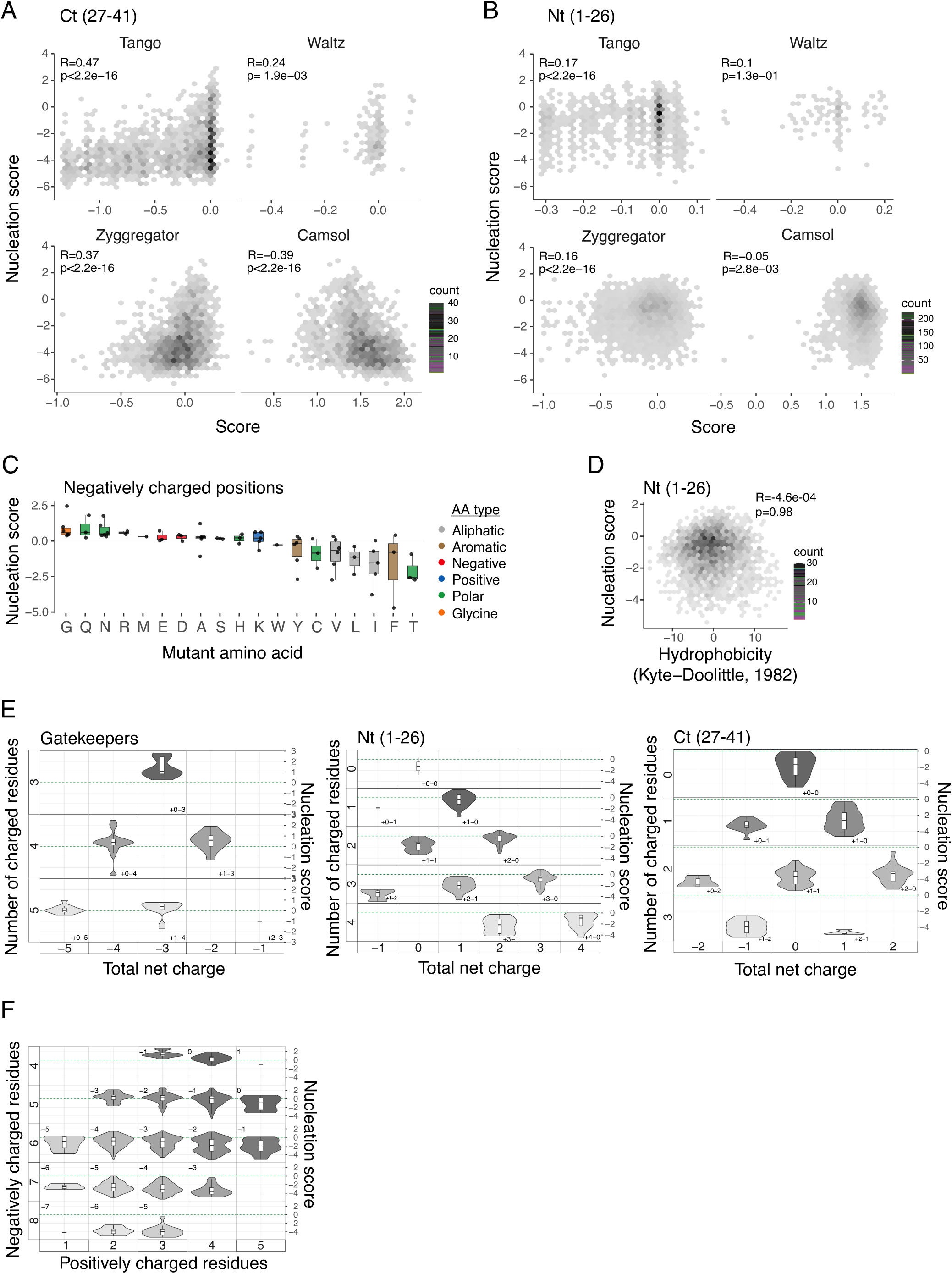
Determinants of Aß nucleation. **A and B** Nucleation scores as a function of aggregation predictors (Tango, Waltz, Zyggregator and Camsol (Tartaglia and Vendruscolo 2008; Fernandez-Escamilla et al. 2004; Oliveberg 2010; Sormanni, Aprile, and Vendruscolo 2015)) for single and double aa mutants in the Ct (27-41) (**A**) and Nt (1-26) (**B**) clusters (gatekeeper positions are excluded from the N-terminal and C-terminal classes). Only double mutants with both mutations in the indicated position-range were used. Weighted Pearson correlation coefficient and p-value are indicated. **C** Effect of each mutant aa on nucleation for the 6 negatively charged positions (D1, E3, D7, E11, E22 and D23). Colour code indicates aa type (see legend). The horizontal line is set at the WT nucleation score (0). **D** Nucleation scores as a function of hydrophobicity changes (Kyte and Doolittle 1982) for single and double aa mutants in the Nt (1-26) cluster. Only double mutants with both mutations in the indicated position-range were used. Weighted Pearson correlation coefficient and p-value are indicated. **E** Nucleation scores distributions arranged by the number of charged residues (y-axis) and the total net charge (x-axis) for single and double aa mutants in the Nt (1-26), Ct (27-41) and gatekeepers (D1, E3, D7, E11, L17, E22 and A42) clusters. Only double mutants with both mutations in the indicated position-range were used. Only polar, charged and glycine aa types were taken into account, for both WT and mutant residues. Colour gradient indicates the total number of charged residues. Numbers inside each cell indicate the number of positive and negative residues. The horizontal line indicates WT nucleation score (0). **F** Nucleation score distributions arranged by the number of negatively charged residues (y-axis) and the number of positively charged (x-axis) for single and double aa mutants in the full peptide (1-42). Only polar, charged and glycine aa types were taken into account, for both WT and mutant residues. Colour gradient indicates the total number of charged residues. Numbers inside each cell indicate the total net charge. The horizontal line indicates WT nucleation score (0). Boxplots represent median values and the lower and upper hinges correspond to the 25th and 75th percentiles respectively. Whiskers extend from the hinge to the largest value no further than 1.5*IQR (interquartile range). Outliers are plotted individually or omitted when the boxplot is plotted together with individual data points or a violin plot.

**figure supplement 4.**
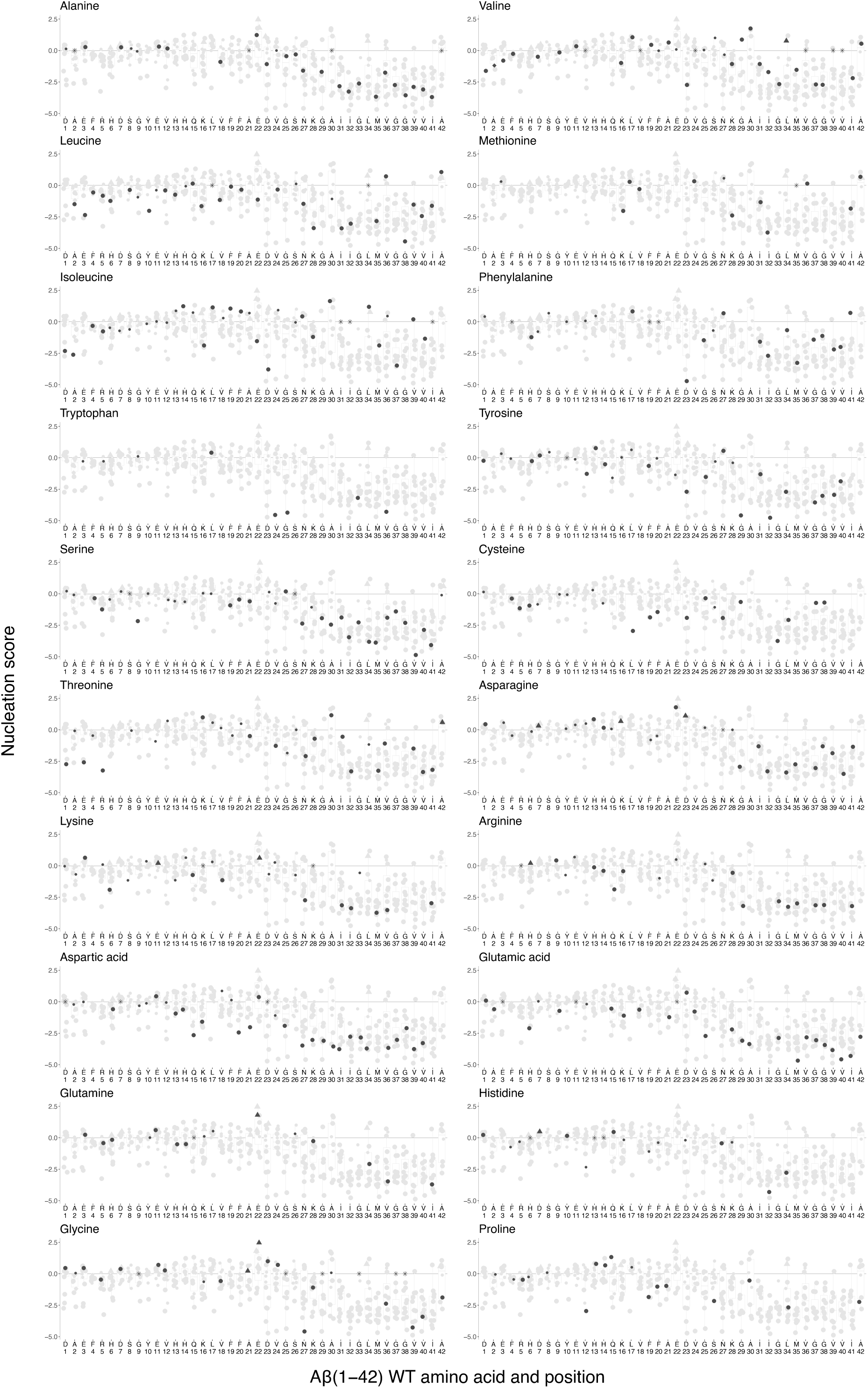
Effect of mutations to each specific aa on Aß nucleation. Mutations to other aa are indicated in grey. The WT AA is indicated with ‘*’. The horizontal line indicates WT nucleation score (0). Point size indicates FDR (big, FDR<0.1; small, FDR>0.1) and shape indicates fAD class (round, VUS; triangle, dominant; rhombus, recessive). Boxplots represent median values and the lower and upper hinges correspond to the 25th and 75th percentiles respectively. Whiskers extend from the hinge to the largest value no further than 1.5*IQR (interquartile range). Outliers are plotted individually or omitted when the boxplot is plotted together with individual data points or a violin plot.

**figure supplement 5.**
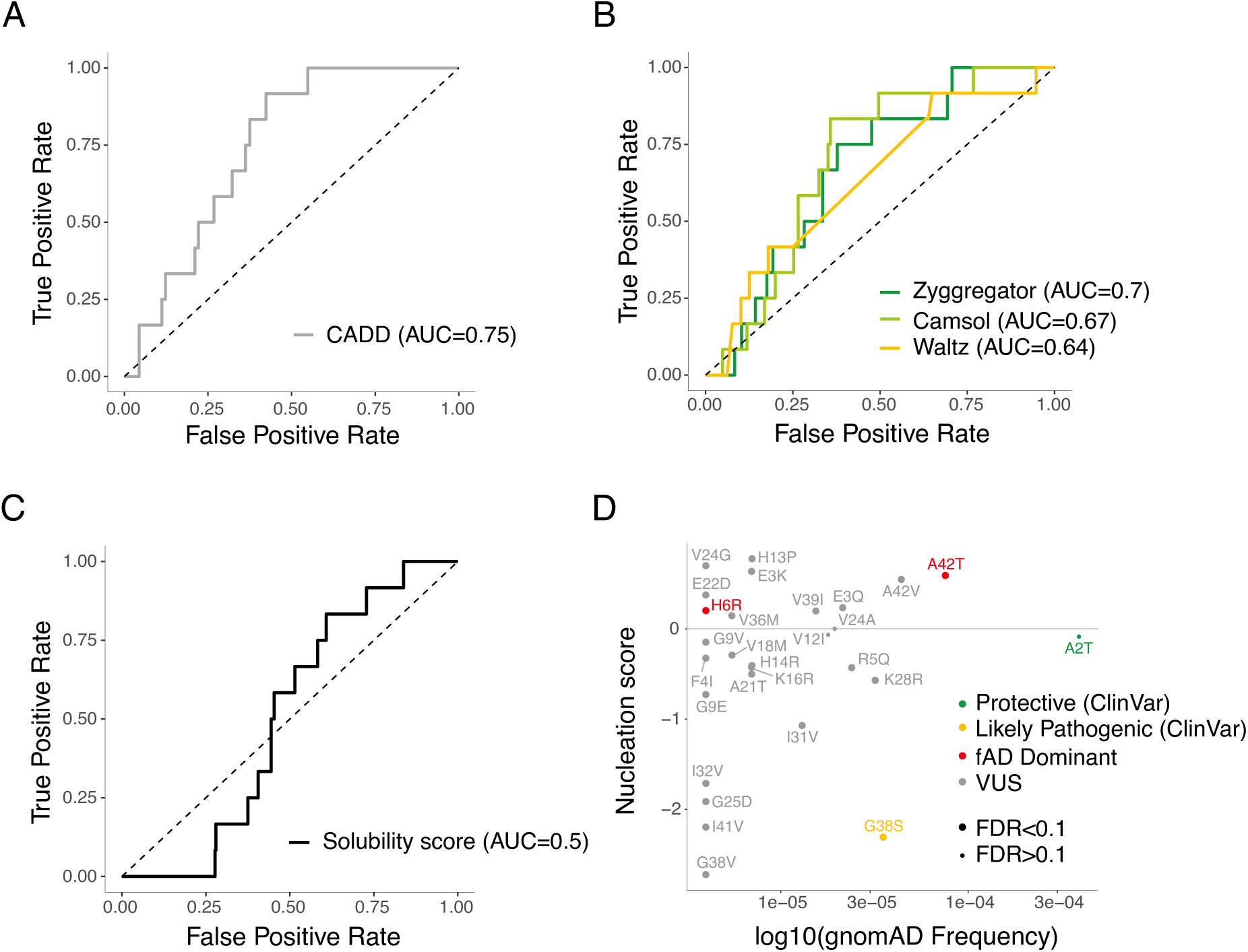
Discrimination of fAD variants by aggregation and variant effect predictors. **A and B** ROC curves built using 12 fAD mutants versus all other single aa mutants in the dataset for variant effect predictors (**A**) and aggregation predictors (**B**). AUC values are indicated in the legend. Diagonal dashed lines indicate the performance of a random classifier. **C** ROC curve built using 12 fAD mutants versus all other single aa mutants available in the referenced study (Gray et al. 2019). AUC value is indicated in the legend. Diagonal dashed line indicates the performance of a random classifier. **D** Comparison between nucleation scores and gnomAD (Karczewski et al. 2020) allele frequencies (https://gnomad.broadinstitute.org/). The horizontal line indicates WT nucleation score (0). The classification of variants is based on Clinvar annotations (https://www.ncbi.nlm.nih.gov/clinvar/).

**figure supplement 6.**
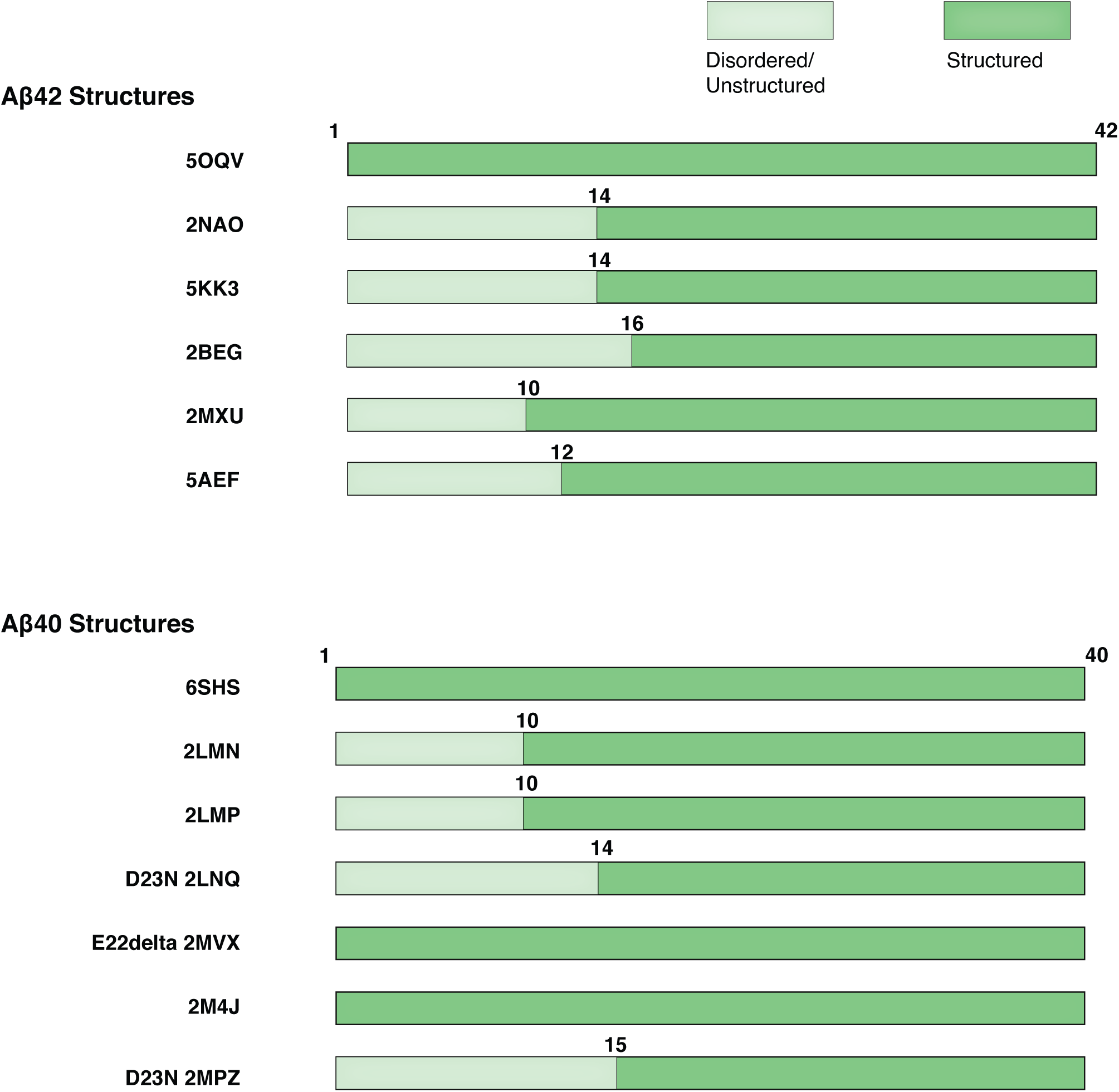
Modular organisation of Aß42 and Aß40 polymorphs. Linear organisation of the Aß42 and Aß40 fibrils. Disordered/unstructured and structured residues are indicated.

**figure supplement 7.**
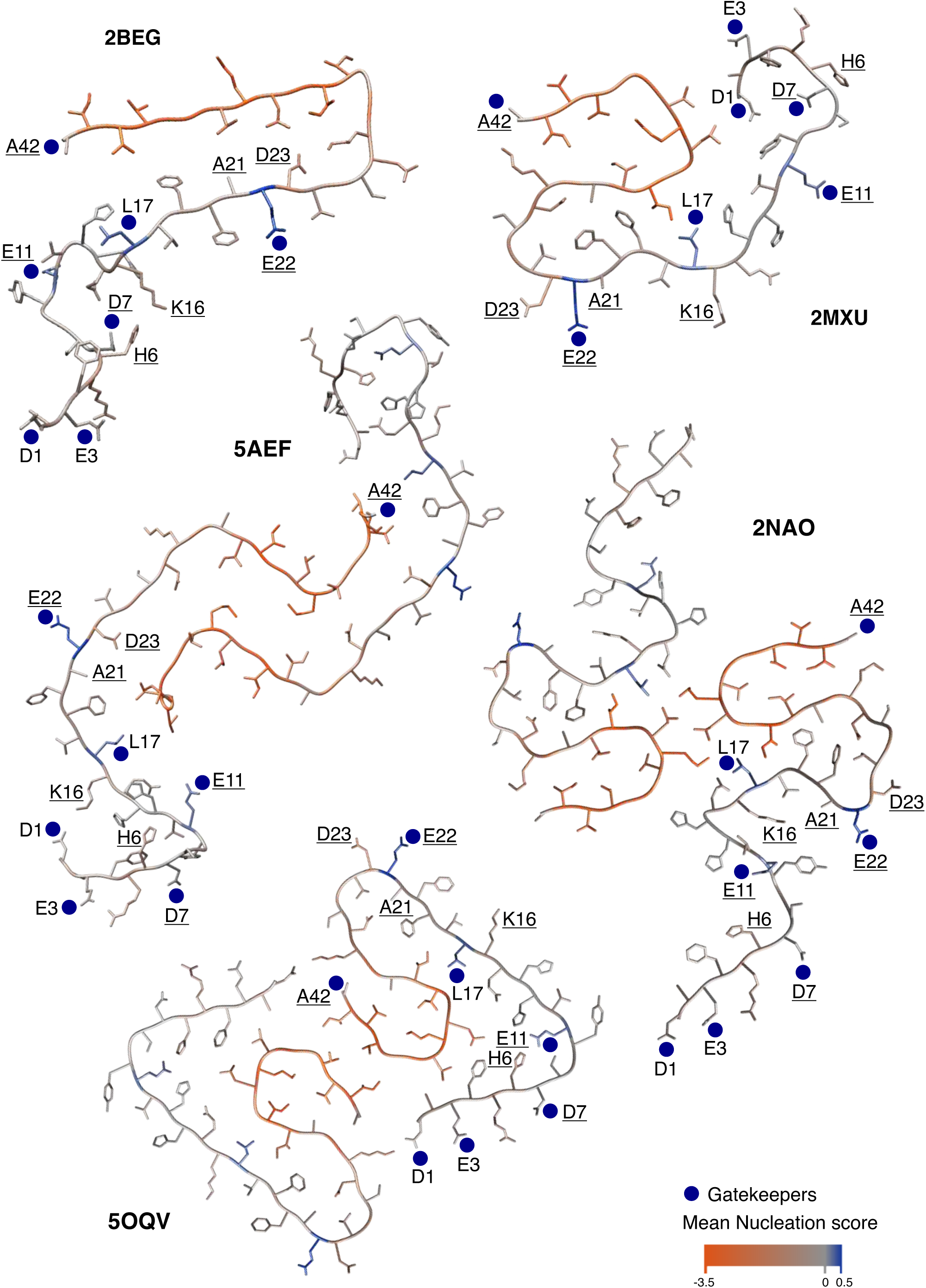
Modular organisation of mutational effects and gatekeepers visualised on Aß42 polymorphs. Average effect of mutations visualised on the cross-section of various Aß amyloid polymorphs: PDB accession 2BEG (Lührs et al. 2005), 2MXU (Xiao et al. 2015), 5AEF (Schmidt et al. 2015), 2NAO (Wälti et al. 2016) and 5OQV (Gremer et al. 2017). Nucleation gatekeeper residues and known fAD mutations positions are indicated by the WT aa identity on one of the two monomers; gatekeepers are indicated with blue dots and fAD are underlined. A single layer of the fibril is shown and the unstructured N-termini are shown with different random coil conformations for the two Aß monomers.

## Materials and Methods

### Plasmid library construction

The plasmid P_CUP1_-Sup35N-Aβ42 used in this study was a kind gift from the Chernoff lab (Chandramowlishwaran et al. 2018).

The Aβ coding sequence and two flanking regions of 52bp and 72bp respectively upstream and downstream of Aβ were amplified (primers MS_01 and MS_02, Supplementary file 2) by error-prone PCR (Mutazyme II DNA polymerase, Agilent). 30 cycles of amplification and 0.01ng of initial template were used to obtain a mutagenesis rate of 16 mutations/kb, according to the manufacturer’s protocol. The product was treated with DpnI (FastDigest, Thermo Scientific) for 2h and purified by column purification (MinElute PCR Purification Kit, Qiagen). The fragment was digested with EcoRI and XbaI restriction enzymes (FastDigest, Thermo Scientific) for 1h at 37C and purified from a 2% agarose gel (QIAquick Gel Extraction Kit, Qiagen). In parallel, the P_CUP1_-Sup35N-Aβ42 plasmid was digested with the same restriction enzymes to remove the WT Aβ sequence, treated with alkaline phosphatase (FastAP, Thermo Scientific) for 1h at 37C to dephosphorylate the 5’ ends, and purified from a 1% agarose gel (QIAquick Gel Extraction Kit, Qiagen).

Mutagenized Aβ was then ligated into the linearized plasmid in a 5:1 ratio (insert:vector) using a ligase treatment (T4, Thermo Scientific) overnight. The reaction was dialysed with a membrane filter (Merck Millipore) for 1h, concentrated 4x and transformed in electrocompetent *E*.*coli* cells (10-beta Electrocompetent, NEB). Cells were recovered in SOC medium and plated on LB with Ampicillin. A total of 4.1 million transformants were estimated, ensuring that each variant of the library was represented more than 10 times. 50ml of overnight *E*.*coli* culture was harvested to purify the Aβ plasmid library with a midi prep (Plasmid Midi Kit, Qiagen). The resulting library contained 29.9% of WT Aβ, 23.8 % of sequences with 1nt change and 21.8 % of sequences with 2 nt changes.

### Large-scale yeast transformation

*Saccharomyces cerevisiae [psi-pin-] (MATa ade1-14 his3 leu2-3,112 lys2 trp1 ura3-52*) strain (also provided by the Chernoff lab) was used in all experiments in this study(Chandramowlishwaran et al. 2018).

Yeast cells were transformed with the Aβ plasmid library starting from an individual colony for each transformation tube. After an overnight pre-growth culture in YPDA medium at 30C, cells were diluted to OD_600_=0.3 in 175ml YPDA and incubated at 30C 200rpm for ∼5h. When cells reached the exponential phase, they were harvested, washed with milliQ and resuspended in sorbitol mixture (100 mM LiOAc, 10mM Tris pH 8, 1mM EDTA, 1M sorbitol). After a 30min incubation at RT, 5ug of plasmid library and 175μl of ssDNA (UltraPure, Thermo Scientific) were added to the cells. PEG mixture (100mM LiOAc, 10mM Tris pH 8, 1mM EDTA pH 8, 40% PEG3350) was also added and cells were incubated for 30min at RT and heat-shocked for 15min at 42C in a water bath. Cells were harvested, washed, resuspended in 350ml recovery medium (YPD, sorbitol 0.5M, 70mg/L adenine) and incubated for 1.5h at 30C 200rpm. After recovery, cells were resuspended in 350ml -URA plasmid selection medium and allowed to grow for 50h. Transformation efficiency was calculated for each tube of transformation by plating an aliquote of cells in -URA plates. Between 1-2.5 million transformants/tube were obtained. Two days after transformation, the culture was diluted to OD_600_=0.02 in 1L -URA medium and grown until the exponential phase. At this stage, cells were harvested and stored at −80C in 25% glycerol.

### Selection experiments

Three independent replicate selection experiments were performed. Tubes were thawed from the −80C glycerol stocks and mixed proportionally to the number of transformants in a 1L total -URA medium at OD_600_=0.05. A minimum of 3.7 million yeast transformants were used for each replicate to ensure the coverage of the full library and reaching therefore a 10x coverage of each variant.

Once the culture reached the exponential phase, cells were resuspended in 1L protein inducing medium (-URA, 20% glucose, 100uM Cu_2_SO_4_) at OD_600_=0.05. As a result, each variant was represented at least 100 times at this stage. After 24h the input pellets were collected by centrifuging 220ml of cells and stored at −20C for later DNA extraction (input pellets). In parallel, 18.5 million cells of the same culture underwent selection, with a starting coverage of at least 50 copies of each variant in the library. For selection, cells were plated on -ADE-URA selective medium in 145cm^2^ plates (Nunc, Thermo Scientific) and let grow for 7 days at 30C. Colonies were then scraped off the plates and recovered with PBS 1x to be centrifuged and stored at −20C for later DNA extraction (output pellets).

### DNA extraction

The input and output pellets (3 replicates, 6 tubes in total) were thawed and resuspended in 2ml extraction buffer (2% Triton-X, 1% SDS, 100mM NaCl, 10mM Tris pH 8, 1mM EDTA pH 8), and underwent two cycles of freezing and thawing in an ethanol-dry ice bath (10 min) and at 62C (10min). Samples were then vortexed together with 1.5ml of phenol:chloroform:isoamyl 25:24:1 and 1.5g of glass beads (sigma). The aqueous phase was recovered by centrifugation and mixed again with 1.5ml phenol:chloroform:isoamyl 25:24:1. DNA precipitation was performed by adding 1:10V of 3M NaOAc and 2.2V of 100% cold ethanol to the aqueous phase and incubating the samples at −20C for 1h. After a centrifugation step, pellets were dried overnight at room temperature.

Pellets were resuspended in 1ml resuspension buffer (10mM Tris pH 8, 1mM EDTA pH 8) and treated with 7.5 μl RNase A (Thermo Scientific) for 30min at 37C. The DNA was finally purified using 75 μl of silica beads (QIAEX II Gel Extraction Kit, Qiagen), washed and eluted in 375μl elution buffer.

DNA concentration in each sample was measured by quantitative PCR, using primers (MS_03 and MS_04, Supplementary file 2) that anneal to the origin of replication site of the plasmid at 58C.

### Sequencing library preparation

The library was prepared for high-throughput sequencing in two rounds of PCR (Q5 High-Fidelity DNA Polymerase, NEB). In PCR1, the Aβ region was amplified for 15 cycles at 68C with frame-shifted primers (MS_05 to MS_18, Supplementary file 2) with homology to Illumina sequencing primers. 300 million of molecules were used for each input or output sample. The products of PCR1 were purified with an ExoSAP-IT treatment (Affymetrix) and a column purification step (QIAquick PCR Purification Kit) and then used as the template of PCR2. This PCR was run for 10 cycles at 62C with Illumina indexed primers (MS_19 to MS_25, Supplementary file 2) specific for each sample (3 inputs and 3 outputs). The 6 samples were then pooled together equimolarly. The final library sample was purified from a 2% agarose gel with silica beads (QIAEX II Gel Extraction Kit, Qiagen). 125bp paired-end sequencing was run on an Illumina HiSeq2500 sequencer at the CRG Genomics Core Facility.

### Data Processing

FastQ files from paired-end sequencing of the Aß library before (‘input’) and after selection (‘output’) were processed using a custom pipeline (https://github.com/lehner-lab/DiMSum). DiMSum is an R package that uses different sequencing processing tools such as FastQC (http://www.bioinformatics.babraham.ac.uk/projects/fastqc/) (for quality assessment), Cutadapt (Martin 2011) (for constant region trimming) and USEARCH (Edgar 2010) (for paired-end read alignment). Sequences were trimmed at 5’and 3’, allowing an error rate of 0.2 (i.e. read pairs were discarded if the constant regions contained more than 20% mismatches relative to the reference sequence). Sequences differing in length from the expected 126bp, or with a Phred base quality score below 30 were discarded. As a result of this processing, around 150 million total reads passed the filtering criteria.

At this stage, unique variants were aggregated and counted using Starcode (https://github.com/gui11aume/starcode). Variants containing indels and nonsynonymous variants with synonymous substitutions in other codons were excluded. The result is a table of variant counts which can be used for further analysis.

For downstream analysis, variants with less than 50 input reads in any of the replicates were excluded and only variants with a maximum of two aa mutations were used.

### Nucleation Scores and error estimates

On the basis of variant counts, the DiMSum pipeline (https://github.com/lehner-lab/DiMSum) was used to calculate nucleation scores (NS) and their error estimates. For each variant in each replicate NSs were calculated as:

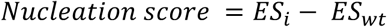

Where *ES*_*i*_ *= log(F*_*i*_ *OUTPUT) − log(F*_*i*_ *INPUT)* for a specific variant and *ES*_*wt*_ *= log(F*_*wt*_ *OUTPUT) − log(F*_*wt*_ *INPUT)* for Aß WT.

DiMSum models measurement error of NS by assuming that variants with similar counts in input and output samples have similar errors. Based on errors expected from Poisson-distributed count data, replicate-specific additive and multiplicative (one each for input and output samples) modifier terms are fit to best describe the observed variance of NS across all variants simultaneously.

After error calculation, NS were merged by using the error-weighted mean of each variant across replicates and centered using the error-weighted means frequency of synonymous substitutions arising from single nt changes. Merged NS and NS for each independent replicate, as well as their associated error estimates, are available in Supplementary file 3.

Nonsense (stop) mutants were excluded for the analysis except when indicated (Figure. 2A and C, figure supplement 2A).

### K-medoids clustering

We used k-medoids, or the partitioning around medoids (PAM) algorithm, to cluster the matrix of single aa variant nucleation scores estimates by residue position with the number of clusters estimated by optimum average silhouette width, for values of K in [1,10]. The silhouette width is a measure of how similar each object (in this case residue position) is to its own cluster. In order to take into account uncertainty in nucleation scores estimates in the determination of the optimum number of clusters, we repeated this analysis after random resampling from the NS (error) distributions of each single aa variant (n=100). Based on this clustering, we defined the N-terminus as aa 1-26 and the C-terminus as aa 27-41. Seven positions where as many (or more) single mutations increase as decrease nucleation were defined as ‘gatekeepers’ (D1, E3, D7, E11, L17, E22, A42) and excluded from the N- and C-terminus classes.

### Amino acid properties, aggregation and variant effect predictors

NSs were correlated with aa properties and scores from aggregation, solubility and variant effect prediction algorithms. Pearson correlations were weighted based on the error terms associated with the NS of each variant using the R package “weights”. The aa property features were retrieved from a curated collection of numerical indices representing various physicochemical and biochemical properties of aas (http://www.genome.jp/aaindex/). We also used a principal component of these aa properties from a previous work (PC1 (Bolognesi et al. 2019)) that relates strongly to changes in hydrophobicity. For each variant (single and double aa mutants), the values of a specific aa property represent the difference between the mutant and the WT scores.

For the aggregation and solubility algorithms (Tango (Fernandez-Escamilla et al. 2004), Zyggregator (Tartaglia and Vendruscolo 2008), CamSol (Sormanni, Aprile, and Vendruscolo 2015) and Waltz (Oliveberg 2010)), individual residue-level scores were summed to obtain a score per aa sequence. We then calculated the log value for each variant relative to the WT score (single and double aa mutants for Tango, Zyggregator, CamSol and single aa mutants for Waltz). For the variant effect predictors (Polyphen (Adzhubei, Jordan, and Sunyaev 2013) and CADD (Rentzsch et al. 2019)) we also calculated the log value for each variant (only single aa mutants) but in this case values were scaled relative to the lowest predicted score.

### fAD, gnomAD & Clinvar variants

The table of fAD mutations used in this study was taken from https://www.alzforum.org/mutations/app. Allele frequencies of APP variants were retrieved from gnomAD (Karczewski et al. 2020) (https://gnomad.broadinstitute.org/) and the clinical significance of variants was taken from their Clinvar (Landrum et al. 2014) classification (https://www.ncbi.nlm.nih.gov/clinvar).

ROC curves were built and AUC values were obtained using the ‘pROC’ R package.

### PDB structures

The coordinates of the following PDB structures were used for Figure 5, figure supplement 6 and 7: 5OQV, 2NAO, 5KK3, 2BEG, 2MXU, 5AEF, 6SHS, 2LMN, 2LMP, 2LNQ, 2MVX, 2M4J, 2MPZ (Gremer et al. 2017; Colvin et al. 2016; Wälti et al. 2016; Lührs et al. 2005; Xiao et al. 2015; Schmidt et al. 2015; Kollmer et al. 2019; Lu et al. 2013; Qiang et al. 2012; Sgourakis, Yau, and Qiang 2015; Schütz et al. 2015).

## Data and code availability

Raw sequencing data and the processed data table (Supplementary file 3) have been deposited in NCBI’s Gene Expression Omnibus (GEO) as record GSE151147. All code used for data analysis is available at https://github.com/BEBlab

## Author Contribution

B.L and B.B designed the study. M.S performed experimental work. M.S, A.J.F and M.B analysed the data. M.S, B.B and B.L wrote the manuscript.

### Acknowledgements

Work in the lab of B.B. is supported by the Spanish Ministry of Science, Innovation and Universities through the project RTI2018-101491-A-I00 (MICIU / FEDER), by the CERCA Program/Generalitat de Catalunya and by funding from the Agencia de Gestio d’Ajuts Universitaris i de Recerca (2019FI_B 01311) to M.S. Work in the lab of B.L. is supported by a European Research Council (ERC) Consolidator grant (616434), the Spanish Ministry of Science, Innovation and Universities (BFU2017-89488-P and SEV-2012-0208), the Bettencourt Schueller Foundation, Agencia de Gestio d’Ajuts Universitaris i de Recerca (AGAUR, 2017 SGR 1322.), and the CERCA Program/Generalitat de Catalunya. We thank the Chernoff lab for kindly providing strains and plasmids and the CRG Genomics core facility for their assistance with sequencing.

## Supplementary Materials

Supplementary file 1 to 3

figure supplement 1 to 7

